# Transcriptional signature in microglia isolated from an Alzheimer’s disease mouse model treated with scanning ultrasound

**DOI:** 10.1101/2021.12.20.473590

**Authors:** Gerhard Leinenga, Liviu-Gabriel Bodea, Jan Schröder, Giuzhi Sun, Yi Chen, Alexandra Grubman, Jose M. Polo, Jürgen Götz

## Abstract

**Rationale:** Intracranial scanning ultrasound combined with intravenously injected microbubbles (SUS^+MB^) has been shown to transiently open the blood-brain barrier and reduce amyloid-β (Aβ) pathology in the APP23 mouse model of Alzheimer’s disease (AD). This has been accomplished, at least in part, through the activation of microglial cells; however, their response to the SUS treatment is only incompletely understood.

**Methods:** Wild-type (WT) and APP23 mice were subjected to SUS^+MB^, using non-SUS^+MB^-treated mice as sham controls. After 48 hours, the APP23 mice were injected with methoxy-XO4 to label Aβ aggregates, followed by microglial isolation into XO4^+^ and XO4^-^ populations using flow cytometry. Both XO4^+^ and XO4^-^ cells were subjected to RNA sequencing and their transcriptome was analyzed through a bioinformatics pipeline.

**Results:** The transcriptomic analysis of the microglial cells revealed a clear segregation depending on genotype (AD model versus WT mice), as well as treatment (SUS^+MB^ versus sham) and Aβ internalization (XO4^+^ versus XO4^-^ microglia). Differential gene expression analysis detected 278 genes that were significantly changed by SUS^+MB^ in the XO4^+^ cells (248 up/30 down) and 242 in XO^-^ cells (225 up/17 down). Not surprisingly given previous findings of increased phagocytosis of plaques following SUS^+MB^, the pathway analysis highlighted that the treatment induced an enrichment in genes related to the phagosome pathway in XO4^+^ microglia; however, when comparing SUS^+MB^ to sham, the analysis revealed an enrichment in genes involved in the cell cycle in both the XO4^+^ and XO4^-^ microglial population.

**Conclusion:** Our data provide a comprehensive analysis of microglia in an AD mouse model subjected to ultrasound treatment as a function of Aβ internalization, one of the defining hallmarks of AD. Several differentially expressed genes are highlighted, pointing to an ultrasound-induced activation of cell cycle mechanisms in microglial cells isolated from APP23 mice treated with SUS^+MB^.

**Graphical abstract:** 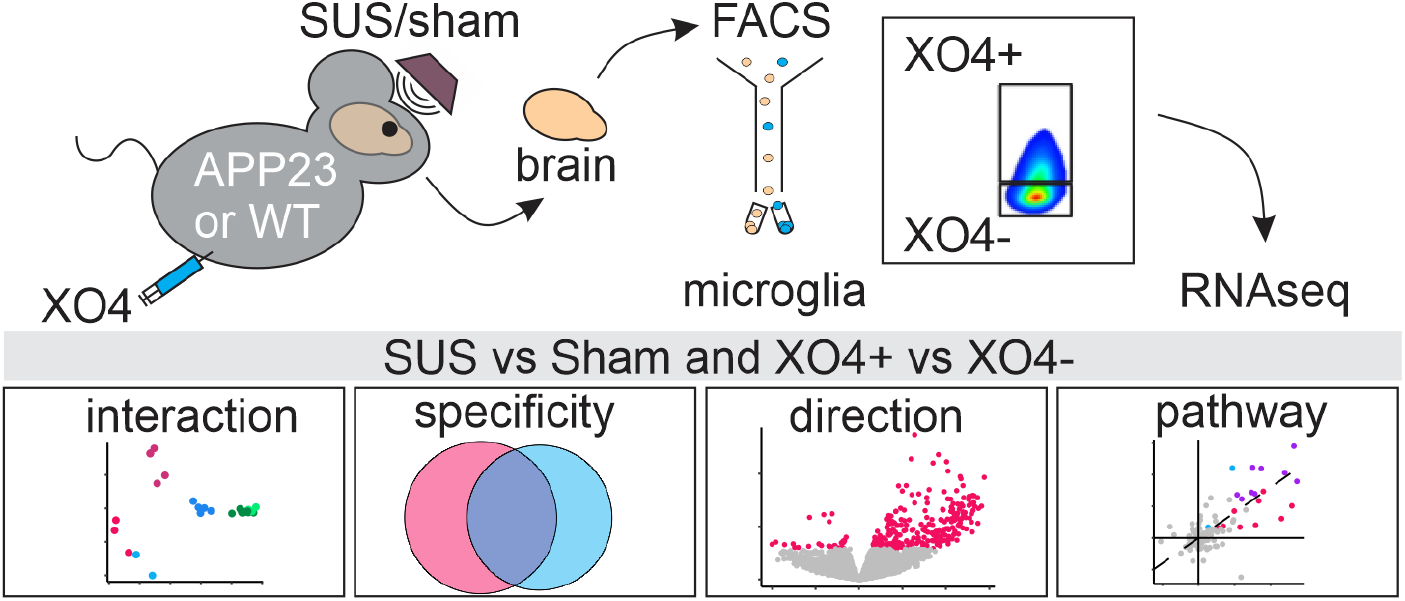

## Introduction

Alzheimer disease (AD) is the most common cause of dementia worldwide. The disease is characterized by progressive and irreversible neurodegeneration. However, given the complexity of the disease combined with a lack of knowledge on how to treat AD efficiently, there is an acute requirement to develop novel treatment strategies [1].

At a histopathological level, AD is characterized by the accumulation of extracellular amyloid-β (Aβ) plaques, intraneuronal tau deposits and increased microglial activation [2]. A broad range of studies have revealed how microglial cells assume both a protective role (through shielding, recognition and removal of Aβ) and a detrimental role (through removal of synapses or release of neurotoxic factors), driving the progression of AD [3]. Transcriptomic studies on microglia have advanced our understanding of the pathogenesis of AD at the level of transcriptional network dynamics, highlighting important molecular players depending on the different phases of the disease [4-6]. Microglia are known to phagocytose aggregated forms of Aβ, and it has been proposed that deficiencies in this process may contribute to late-onset AD [7] and metabolic labelling in humans indicated that clearance of Aβ is impaired in AD [8]. Recently, it has been shown that microglia that have taken up amyloid differ in their transcriptional signature in comparison with microglia that do not contain amyloid [9].

An obstacle to treating AD is the blood-brain barrier which prevents large molecules such as antibodies from entering the brain, with IgG having 0.1% transfer across the blood-brain-barrier (BBB) [10]. Approaches to modify anti-Aβ antibodies to increase levels in the brain are in development [11] along with other approaches to circumvent the BBB.

Studies in animal models of AD have indicated that repeated transient blood-brain barrier (BBB) openings that are achieved throughout the entire brain using intracranial ultrasound in a scanning mode together with intravenously injected microbubbles (SUS^+MB^) significantly clear amyloid plaques. One study reported that plaque reduction can occur as fast as 48 hours after BBB opening [12], and we have shown that this process occurs through microglial phagocytosis [13]. Ultrasound-mediated bioeffects including microglial activation have also been demonstrated by specifically targeting the hippocampus [14, 15], but the therapeutic benefit seems to be most pronounced when the brain is treated more globally [13]. Of note, this clearing process requires BBB opening [16] and is even effective at reducing Aβ pathology in 22-month-old senescent mice [17]. Combination treatments with ultrasound for delivery of anti-Aβ antibodies such as Aducanumab that has been recently approved by the FDA [18], or an anti-pyroglutamylated Aβ antibody [19], led to more effective plaque removal and behavioral improvements than in those observed in mice that were treated with either ultrasound alone or antibodies alone.

Ultrasound-mediated BBB opening has also been achieved in a small safety trial that revealed tolerability in patients with mild AD when a small region of the frontal cortex was targeted [20]. A subsequent study found that the BBB could be opened in parts of the hippocampus [21], with a modest reduction in the amyloid PET signal following three treatments with ultrasound over a 6-month period [22]. In all these studies, BBB opening by ultrasound was shown to be safe and reversible in that the BBB was fully restored after 24 hours.

Several mechanisms have been proposed to explain how BBB opening leads to amyloid plaque reduction, including the uptake of endogenous immunoglobulins [23] or albumin binding to amyloid [13], followed by microglial phagocytosis of Aβ and lysosomal digestion. Here, to gain a better understanding of how the combination of SUS treatment and Aβ internalization affects microglial cells, we analyzed the transcriptional profile of microglia isolated from APP23 mice (a model of AD) that had been subjected to SUS^+MB^. By using a fluorescent dye to detect Aβ internalization within the microglia we identified differences between the microglial cells from mice treated with or without ultrasound, as well as between cells that had internalized Aβ or not.

## Materials and methods

### Animals

In this study, we have used APP23 mice (harboring the AD Swedish K670M, N671L double mutation in the APP gene [24]) and wild-type (WT) mice. The animals were maintained on a 12 h light/dark cycle and housed in a PC2 facility with *ad libitum* access to food and water. All experimental procedures in this study (**Fig 1A**) were approved by the University of Queensland Animal Ethics Committee (AEC) (QBI/412/14/NHMRC and QBI/554/17/NHMRC), and Monash University AEC (17241) and were conducted in compliance with the ARRIVE guidelines (Animal Research: Reporting in Vivo Experiments).

**Figure 1.**
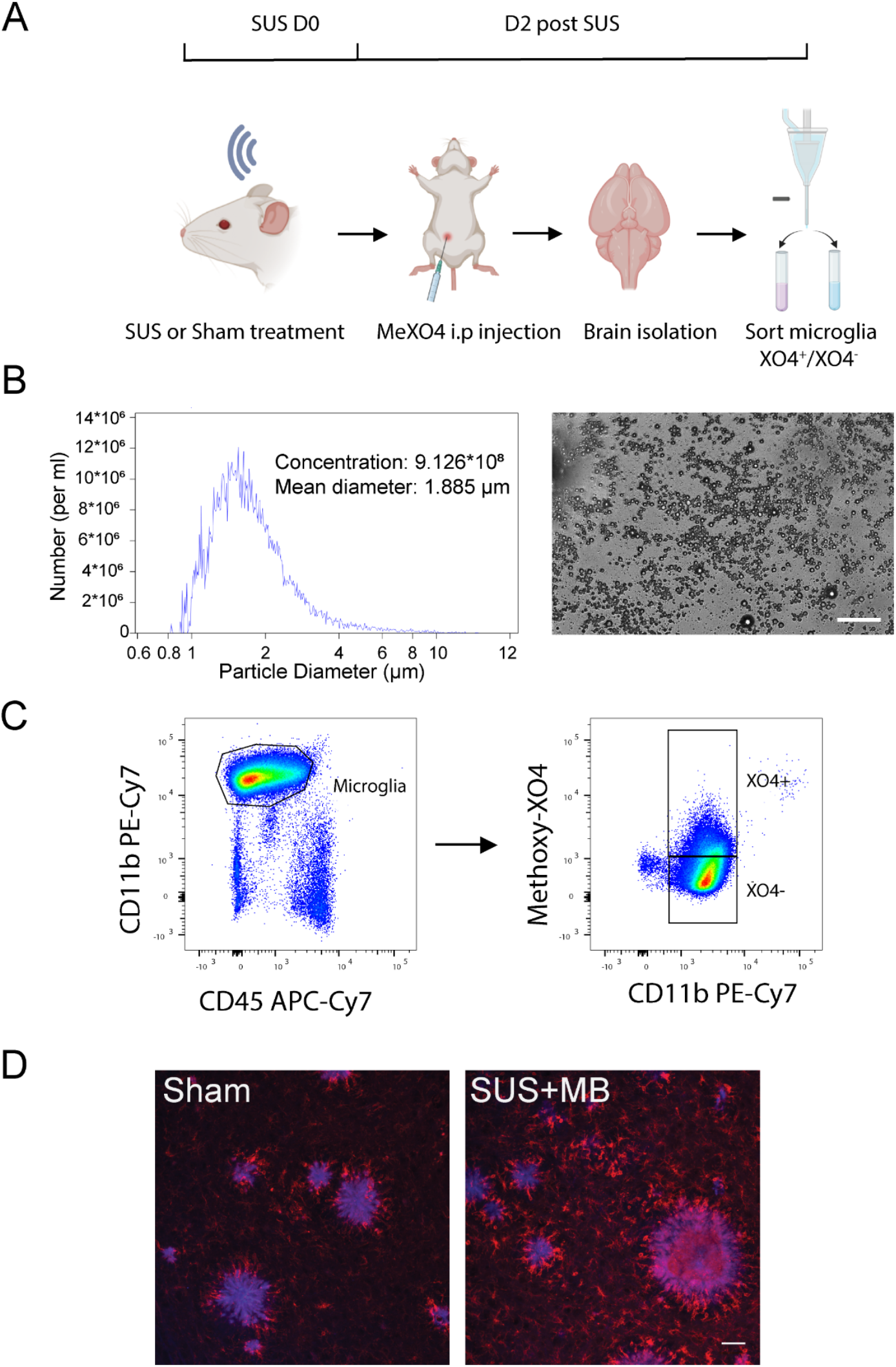
Experimental design and gating strategy to isolate XO4^+^ and XO4^-^ microglia. **(A)** Scanning ultrasound (SUS^+MB^) or sham (no ultrasound) treatment was applied to APP23 transgenic and wild-type (WT) mice. Two days post-treatment, the mice received a single injection with methoxy-XO4 that binds Aβ two hours before euthanasia and collection of brain tissue. The brains of the mice were harvested and lysed to form a single-cell suspension, followed by FACS-based isolation of XO4^+^ and XO4^-^ microglial cells. (**B)** In-house prepared microbubbles were used for scanning ultrasound (SUS^+MB^) and their size and concentration were measured by Coulter Counter. **(C)** The gating strategy used to isolate microglial cells into XO4^+^ and XO4^-^ populations via FACS using CD11b and CD45 antibodies to isolate a pure population of microglia, and Methoxy-XO4 fluorescence to isolate microglia cells that contain methoxy-XO4 bound to Aβ. **(D)** Methoxy-XO4 binds to Aβ plaques in the brains of APP23 mice (blue) in close proximity to Iba1-positive microglia (red). Scale bar 50 μm.

### SUS treatment

Microbubbles comprising a phospholipid shell and octafluoropropane gas core were prepared in-house. 1,2-distearoyl-sn-glycero-3-phosphocholine (DSPC) and 1,2-distearoyl-sn-glycero-3-phosphoethanolamine-N-[amino(polyethylene glycol)-2000] (DSPE-PEG2000) (Avanti Polar Lipids) were mixed in a 9:1 molar ratio and dissolved in chloroform (Sigma), after which the chloroform solvent was evaporated under vacuum. The dried phospholipid cake was then dissolved in PBS with 10% glycerol to a concentration of 1 mg lipid/ml and heated to 55°C in a sonicating water bath. The solution was placed in a 1.5 ml glass high-performance liquid chromatography (HPLC) vial with the air in the vial replaced with octafluoropropane gas (Arcadophta). The microbubbles were activated on the day of the experiment by agitation of the vial in a dental amalgamator at 4,000 rpm for 45s. Activated microbubbles were measured with a Multisizer 4e coulter counter which reported a mean diameter of 1.885 μm and a concentration of 9.12×10^8^ microbubbles/ml. These microbubbles were also observed to be polydisperse under a microscope (**Fig 1B**).

For treatment delivery, an integrated focused ultrasound system (Therapy Imaging Probe System, TIPS, Philips Research) was used. This system consisted of an annular array transducer with a natural focus of 80 mm, a radius of curvature of 80 mm, a spherical shell of 80 mm with a central opening of 31 mm diameter, a 3D positioning system, and a programmable motorized system to move the ultrasound focus in the x and y planes to cover the entire brain area. A coupler mounted to the transducer was filled with degassed water and placed on the head of the mouse with ultrasound gel for coupling, to ensure unobstructed propagation of the ultrasound to the brain.

For SUS^+MB^ applications, mice were anesthetized with ketamine (90 mg/kg) and xylazine (6 mg/kg) and the hair on their head was shaved and depilated. They were then injected retro-orbitally with 1 μl/g body weight of microbubble solution and placed under the ultrasound transducer with the head immobilized. A heating pad was used to maintain body temperature. Parameters for the ultrasound delivery were 1 MHz center frequency, 0.65 MPa peak negative pressure, 10 Hz pulse repetition frequency, 10% duty cycle, and a 6 second sonication time per spot. The focus of the transducer was 1.5 mm x 12 mm in the transverse and axial planes, respectively. The motorized positioning system moved the focus of the transducer array in a grid with 1.5 mm spacing between individual sites of sonication so that ultrasound was delivered sequentially to the entire brain as described previously [13, 18]. Mice typically received a total of 24 spots of sonication in a 6×4 raster grid pattern. For the sham treatment, mice received all injections and were placed under the ultrasound transducer, but no ultrasound was emitted. The time between injecting microbubbles and commencing ultrasound delivery was 60 ±10 s and the duration of sonication was approximately 3 min (total time from microbubble injection approximately 4 min).

### Acute isolation of microglia and fluorescence activated cell sorting

Two hours prior to brain harvest, mice were injected intraperitoneally with methoxy–X04 (2 mg/ml in 1:1 ratio of DMSO to 0.9% (w/v) NaCl, pH 12) at 5 mg/kg. Mice were euthanized by CO_2_ and transcardially perfused with ice-cold PBS prior to brain extraction. Whole brains, excluding the brain stem, olfactory bulbs and cerebellum, were dissected for microglial isolation. Single cell suspensions were prepared by mechanical dissociation using meshes of decreasing sizes from 250 μm to 70 μm and suspensions were enriched for microglia by density gradient separation. Briefly, the cell pellet was resuspended in 70 % (v/v) isotonic Percoll (1x PBS + 90 % (v/v) Percoll), overlayed with 37 % (v/v) isotonic Percoll and centrifuged with slow acceleration and no brake at 2,000 g for 20 min at 4 °C. The microglia-enriched cell population isolated from the 37 %-70 % interphase was diluted 1:5 in ice-cold PBS and recovered by cold centrifugation at maximum speed for 1 min in microcentrifuge tubes. The cell pellet was then stained with antibodies to microglial cell surface markers (CD11b-PE Cy7, 1:200 Biolegend, # 101216; CD45-APC Cy7, 1:200, BD Biosciences # 103116;) for isolation using the FACSAria™ III cell sorter (**Fig 1C**). For mice without endogenous expression of CX3CR1-GFP, CX3CR1 was stained using CX3CR1-FITC (1:100, Biolegend, #149019). Microglia were defined as live/propidium iodide (PI)^−^ (Sigma-Aldrich, St. Louis, MO, #P4864), CD11b^+^, CD45^low^, CX3CR1^+^ cells. The XO4^+^ population gate was set using methoxy-XO4-injected wild-type animals. X04^+^ and X04^−^ microglial populations were sorted separately for further analysis.

### Immunohistology

APP23 mice were treated with SUS^+MB^ and were perfused with PBS and drop fixed in 4% paraformaldehyde in PBS. They were then cryoprotected in 30% sucrose in PBS and sectioned at 40 μm with a freezing sliding microtome. Microglia were immunostained with anti-Iba1 antibody (Wako, JP 1:1,000) followed by incubation with an anti-rabbit secondary antibody AlexaFluor 568 conjugate (Invitrogen). Sections were then co-stained with 10 μM methoxy-XO4 (Tocris Bioscience) in PBS with 20% ethanol before cover-slipping. Images were obtained with a spinning disk confocal microscope (Nikon Diskovery) with a 20x objective, acquiring z-stacks through the entire depth of the section (**Fig 1D**).

### Bulk RNA-seq and bioinformatics analysis

RNA extraction from FACS-sorted microglia was performed using the RNeasy Micro Kit (Qiagen, #74004) and RNA quality was assessed using the Bioanalyser (Agilent RNA 6000 Pico kit; #5067-1513). The libraries were prepared using microglia RNA samples with RIN value ≥ 7.9 as previously described [9]. Sequencing reads were mapped to the mouse transcriptome reference genome (GRCm38) using STAR (v020201). We then established read counts for each gene using featureCounts (v1.5.2). The bulk RNA-seq read counts were further analysed using R (v4.0.2), limma (v3.42.2), and edgeR (v3.28.1). Data handling and plotting were performed using tidyverse (v1.3.0). In detail, we first removed lowly or non-expressed genes with the *filterByExpr* function, and we calculated TMM (trimmed mean of m-values) normalisation factors to remove composition bias using *calcNormFactors*. To visualise dimensionality reduction of the sequencing data, we first removed unwanted variation in the data with *removeBatchEffect*. The principal component analysis (PCA plot, **Fig. 2A**) is using *prcomp* and *ggplot* to visualise the remaining variance in the data. The heatmap (**Fig. 2B**) is plotting the DEGs from either contrast using the wardD2 method and the *pheatmap* function. To determine differentially expressed genes (DEGs), we first determined the gene-wise variance trends using *voom*. Then, we built a linear model using the *lmFit* function and all combinations of genotype and treatment, batch, and sex as covariates. The Venn diagrams (**Fig. 3**) are generated with the VennDiagram package (v1.7.0). **Tables 1,2** show the top 50 DEGs for the two contrasts of interest (ranked by absolute log fold change). Pathway and gene ontology enrichments are calculated from the respective DEG lists using the *kegga* and *goana* functions **(Tables 3**,**4)**.

**Figure 2.**
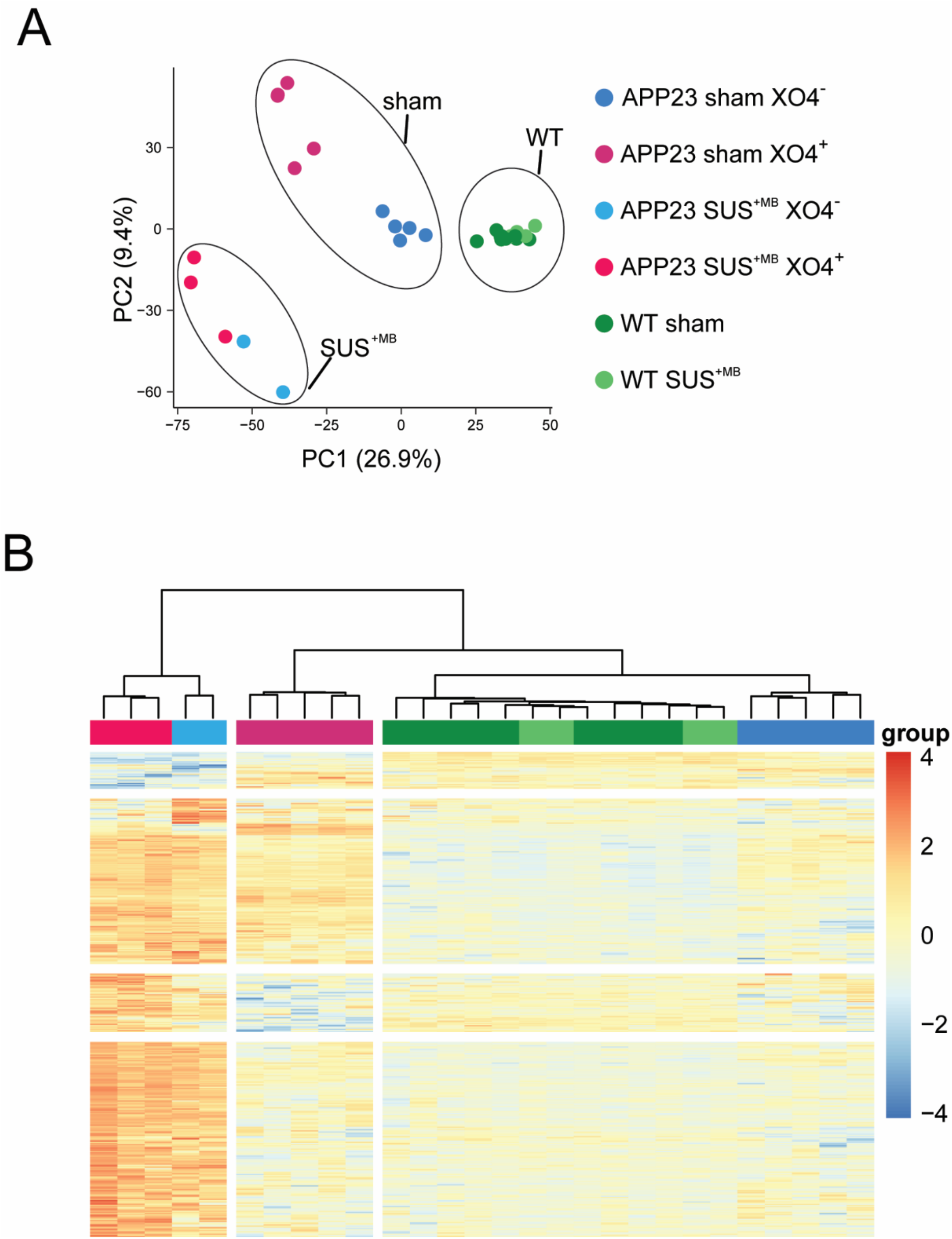
Cluster analysis distinguishes between genotype-, treatment-, and Aβ-dependent microglial phenotypes. **(A)** Principal component analysis reveals the presence of a WT microglia cluster independent of SUS^+MB^ treatment that is segregated from cells of APP23 origin (further clustered dependent on both the SUS^+MB^ treatment and their Aβ content) Ovals indicate treatment groups: SUS^+MB^, Sham, and WT mice. (**B)** Hierarchical clustering of the differential genes (SUS^+MB^ versus sham) reveals a clear segregation between samples of APP23 or WT origin, SUS^+MB^- or sham-treated, and containing Aβ or not.

**Figure 3.**
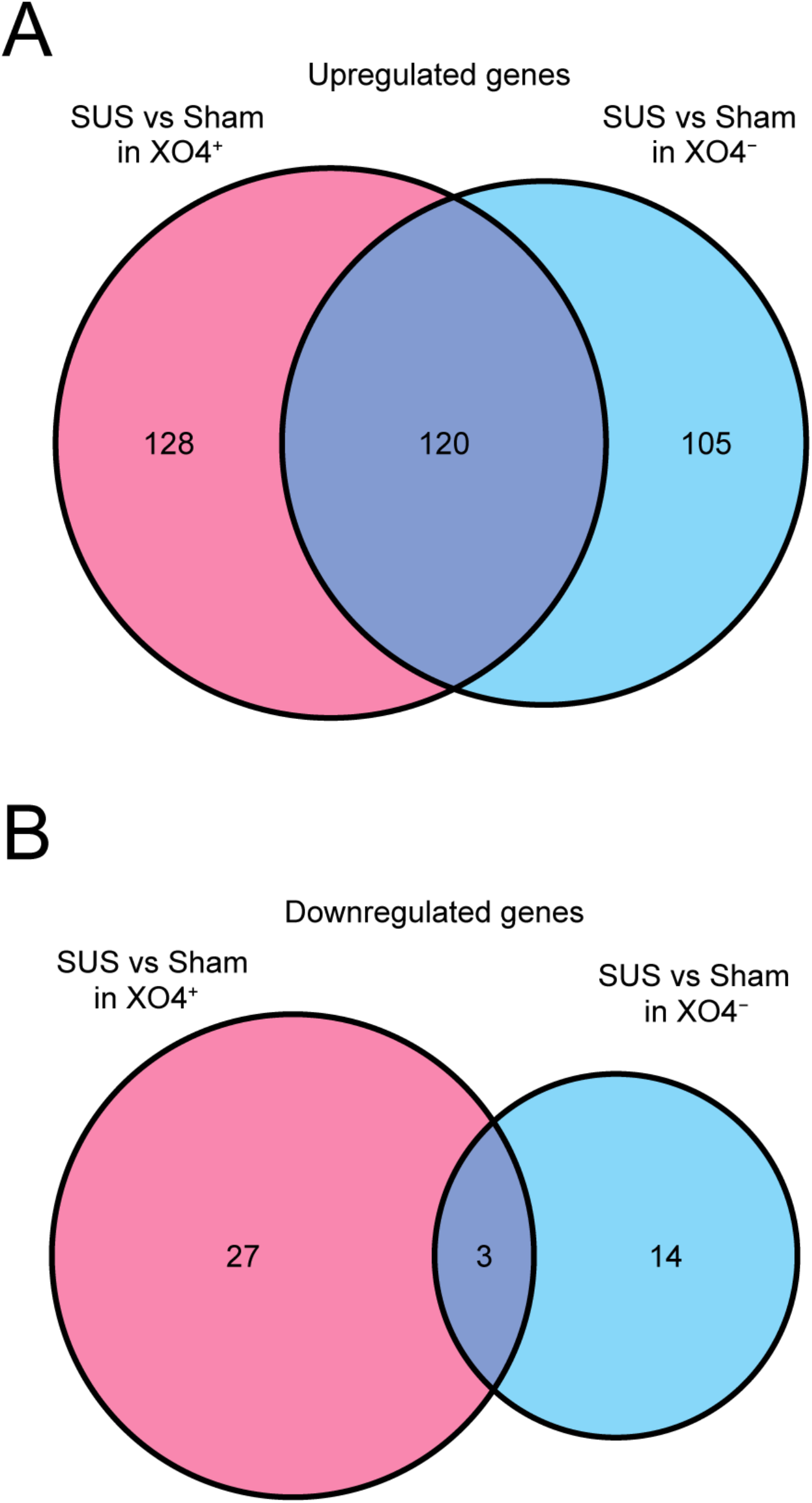
SUS^+MB^ treatment leads to a marked increase in the number of upregulated genes in both XO4^+^ and XO4^-^ microglia, when compared with sham-treated APP23 mice. **(A)** A Venn diagram depicting the number of genes upregulated by SUS^+MB^ distribute similarly between XO4^+^ and XO4^-^ cells, with many genes up in both. (**B**) A larger number of genes were down-regulated in the XO4^+^ cells compared with XO4^-^ cells following SUS^+MB^, with few genes down in both.

**Table 1.** Top 50 dysregulated genes in XO4^+^ microglia from SUS^+MB^ vs sham-treated APP23 mice.

| Gene symbol | logFC | AvgExp | t | Pval | adj PVal | B |
| --- | --- | --- | --- | --- | --- | --- |
| Birc5 | 5.76502 | 1.7672889 | 9.02935 | 2.22E-10 | 3.14E-07 | 13.48506 |
| Cdkn3 | 5.64183 | 1.5432145 | 6.744088 | 1.18E-07 | 3.49E-05 | 7.515466 |
| Kif11 | 5.49926 | 0.2461131 | 6.576116 | 1.91E-07 | 5.14E-05 | 6.97426 |
| Ncapg | 5.47342 | -0.1786981 | 5.464827 | 4.88E-06 | 5.93E-04 | 3.988828 |
| Cdc20 | 5.45743 | 1.0739871 | 7.403038 | 1.82E-08 | 8.52E-06 | 9.133173 |
| Nuf2 | 5.32476 | -0.3289444 | 6.736156 | 1.21E-07 | 3.49E-05 | 7.284114 |
| Ccnb2 | 5.2893 | 1.9141487 | 7.553196 | 1.19E-08 | 6.74E-06 | 9.804982 |
| Aurkb | 5.28106 | 0.1698159 | 7.026936 | 5.26E-08 | 2.05E-05 | 8.075848 |
| Cit | 5.24623 | 0.3082589 | 6.415315 | 3.04E-07 | 7.16E-05 | 6.501784 |
| Hmmr | 5.22306 | 1.186993 | 7.569951 | 1.14E-08 | 6.74E-06 | 9.797002 |
| Esco2 | 5.20983 | 1.0709507 | 4.81764 | 3.25E-05 | 2.80E-03 | 2.353105 |
| Melk | 5.17911 | -0.554891 | 6.172445 | 6.16E-07 | 1.18E-04 | 5.824797 |
| Cdc25c | 5.17482 | -0.7504202 | 5.556821 | 3.73E-06 | 4.68E-04 | 4.188965 |
| Kif23 | 5.13702 | 0.6650897 | 6.945323 | 6.63E-08 | 2.30E-05 | 7.956364 |
| Nek2 | 5.09204 | -0.8391415 | 5.851377 | 1.57E-06 | 2.54E-04 | 4.972077 |
| Troap | 5.07497 | -1.0475974 | 7.38968 | 1.89E-08 | 8.52E-06 | 8.82398 |
| Dlgap5 | 5.0637 | -0.3794831 | 6.798489 | 1.01E-07 | 3.16E-05 | 7.445236 |
| Tk1 | 5.06171 | 1.4926731 | 7.504487 | 1.37E-08 | 7.36E-06 | 9.280781 |
| Mcm10 | 5.04637 | -1.0895909 | 7.260236 | 2.71E-08 | 1.10E-05 | 8.355957 |
| Saa3 | 4.93308 | -1.1465763 | 5.045849 | 1.67E-05 | 1.68E-03 | 2.844773 |
| Plac8 | 4.9074 | 1.316802 | 5.889764 | 1.40E-06 | 2.33E-04 | 5.218269 |
| Clspn | 4.87938 | 1.3914605 | 6.258953 | 4.79E-07 | 9.81E-05 | 6.258822 |
| Mki67 | 4.81598 | 2.6575306 | 7.401992 | 1.82E-08 | 8.52E-06 | 9.451222 |
| Cenpk | 4.81418 | 1.0168275 | 6.273493 | 4.59E-07 | 9.60E-05 | 6.240974 |
| Casc5 | 4.79524 | 1.9047316 | 6.021465 | 9.56E-07 | 1.71E-04 | 5.65795 |
| Ccnb1 | 4.78376 | 0.8942972 | 8.35344 | 1.33E-09 | 1.33E-06 | 11.79537 |
| Ndc80 | 4.77516 | 0.9359373 | 6.74738 | 1.17E-07 | 3.49E-05 | 7.549301 |
| Aspm | 4.77431 | 0.7784471 | 6.30262 | 4.22E-07 | 8.98E-05 | 6.4019 |
| Mns1 | 4.77059 | -0.9456452 | 5.116901 | 1.35E-05 | 1.40E-03 | 2.98948 |
| Stil | 4.71375 | -0.4286108 | 6.896241 | 7.63E-08 | 2.46E-05 | 7.747276 |
| Ankle1 | 4.70824 | -0.8978755 | 5.791278 | 1.87E-06 | 2.91E-04 | 4.711205 |
| Spc24 | 4.69279 | 1.3423116 | 8.911432 | 3.03E-10 | 3.80E-07 | 12.94538 |
| Gpsm2 | 4.65782 | -0.7681863 | 5.503262 | 4.36E-06 | 5.35E-04 | 4.048151 |
| Prc1 | 4.6556 | 2.8666675 | 9.092631 | 1.89E-10 | 3.04E-07 | 13.79044 |
| Cenpi | 4.63601 | 0.7782963 | 5.284361 | 8.29E-06 | 9.00E-04 | 3.53967 |
| Rab13 | 4.6245 | -0.436683 | 5.777597 | 1.95E-06 | 2.91E-04 | 4.733482 |
| Cep55 | 4.61759 | 0.1648057 | 6.537217 | 2.14E-07 | 5.62E-05 | 6.96793 |
| Hist1h2ae | 4.61674 | 0.6086809 | 5.236377 | 9.54E-06 | 1.03E-03 | 3.448414 |
| Cdc6 | 4.59923 | -0.7363785 | 5.380778 | 6.25E-06 | 7.20E-04 | 3.650384 |
| Foxm1 | 4.57058 | 0.2001768 | 4.66003 | 5.15E-05 | 4.18E-03 | 1.887387 |
| Sgol2a | 4.56722 | 0.8256839 | 5.301333 | 7.89E-06 | 8.65E-04 | 3.661214 |
| Tpx2 | 4.5441 | 1.3968306 | 6.899334 | 7.56E-08 | 2.46E-05 | 8.057218 |
| Shcbp1 | 4.5333 | -0.2503641 | 6.219989 | 5.36E-07 | 1.06E-04 | 6.048152 |
| Chaf1a | 4.5075 | -0.3486608 | 4.197819 | 1.95E-04 | 1.29E-02 | 0.699981 |
| Sgol1 | 4.50239 | 1.1036653 | 6.068962 | 8.32E-07 | 1.54E-04 | 5.751895 |
| Ckap2l | 4.49338 | 2.7278769 | 7.949218 | 3.99E-09 | 3.22E-06 | 10.8567 |
| Rad51ap1 | 4.43737 | 1.7528958 | 5.761704 | 2.04E-06 | 3.00E-04 | 4.897387 |
| C330027C09Rik | 4.41274 | 1.2903056 | 4.872646 | 2.77E-05 | 2.50E-03 | 2.500517 |
| Ccnf | 4.38312 | 0.9642677 | 5.68638 | 2.55E-06 | 3.43E-04 | 4.605822 |
| Ckap2 | 4.37671 | 0.4264924 | 5.925534 | 1.26E-06 | 2.20E-04 | 5.33635 |

**Table 2.**
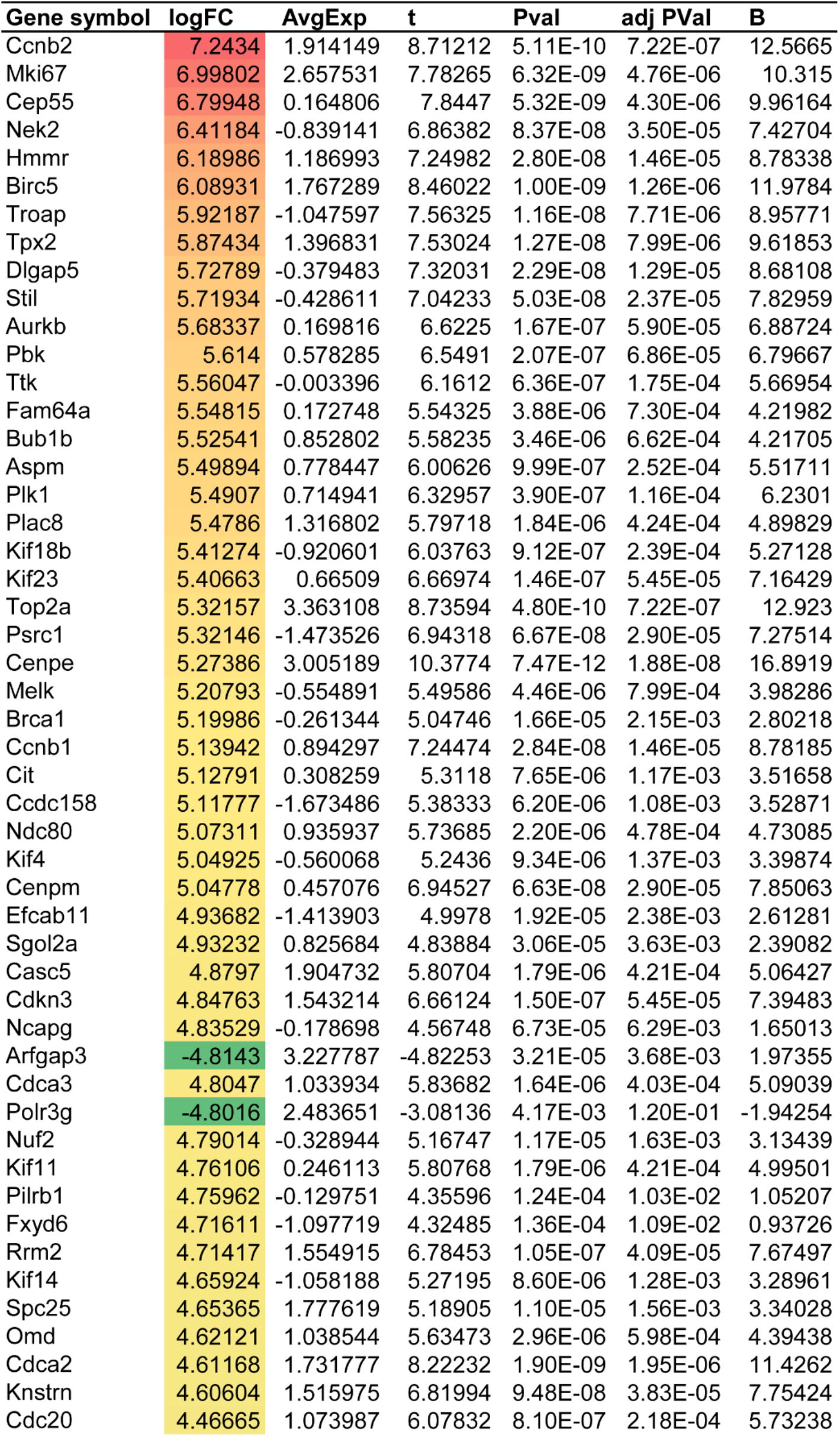
Top 50 dysregulated genes in XO4^-^ microglia from SUS^+MB^ vs sham-treated APP23 mice.

**Table 3.**
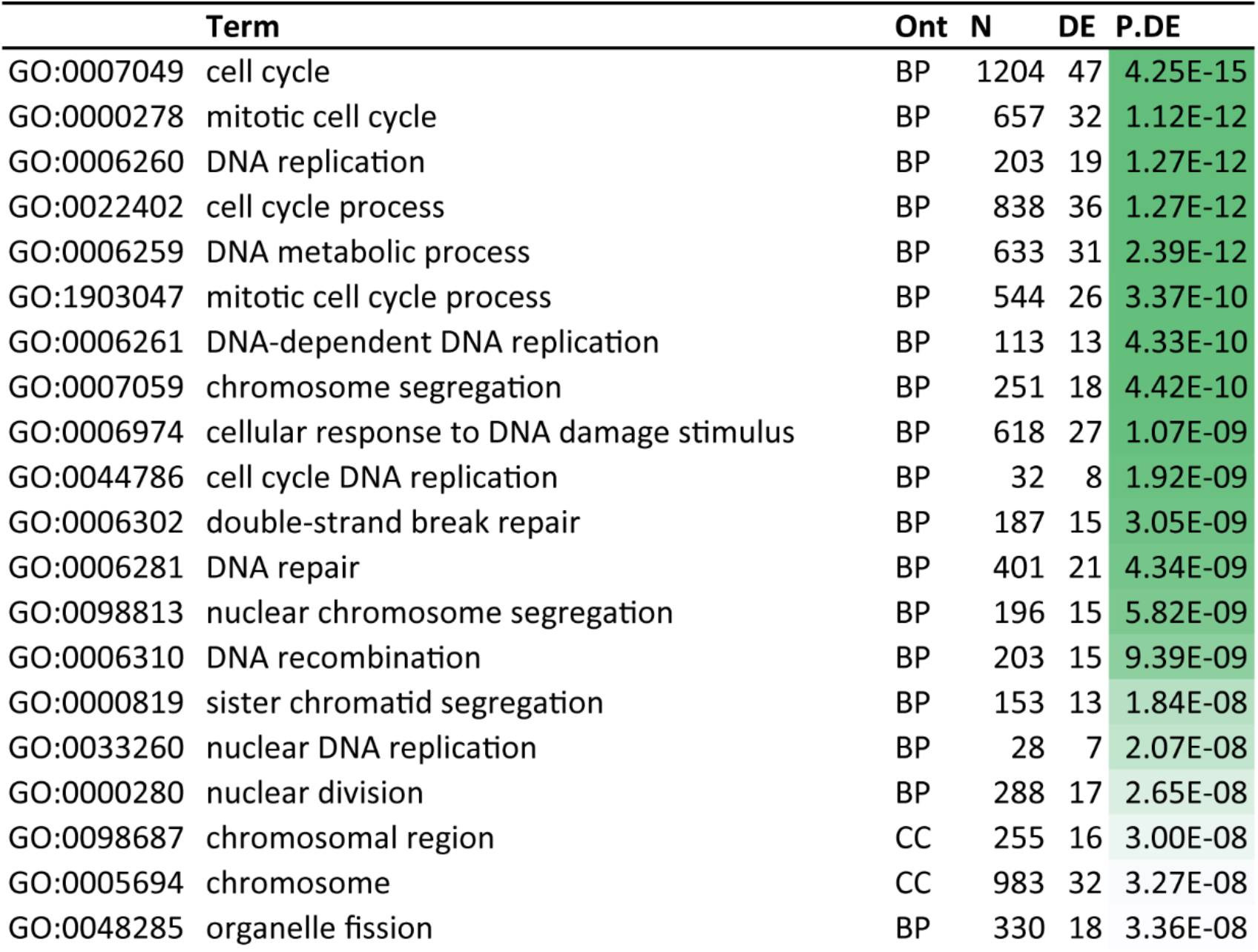
GO pathway enrichment in microglia from SUS^+MB^-versus sham-treated APP23 mice.

**Table 4.** KEGG pathway enrichment in microglia from SUS^+MB^-versus sham-treated APP23 mice.

|  | Pathway | N | DE | P.DE |
| --- | --- | --- | --- | --- |
| path:mmu03030 | DNA replication | 34 | 6 | 2.13E-06 |
| path:mmu04110 | Cell cycle | 115 | 9 | 6.69E-06 |
| path:mmu00670 | One carbon pool by folate | 17 | 4 | 3.55E-05 |
| path:mmu04114 | Oocyte meiosis | 90 | 5 | 3.76E-03 |
| path:mmu05150 | Staphylococcus aureus infection | 28 | 3 | 3.95E-03 |
| path:mmu04145 | Phagosome | 110 | 5 | 8.73E-03 |
| path:mmu04914 | Progesterone-mediated oocyte maturation | 72 | 4 | 9.42E-03 |
| path:mmu00240 | Pyrimidine metabolism | 39 | 3 | 1.01E-02 |
| path:mmu03013 | Nucleocytoplasmic transport | 96 | 4 | 2.47E-02 |
| path:mmu01523 | Antifolate resistance | 25 | 2 | 3.32E-02 |
| path:mmu03008 | Ribosome biogenesis in eukaryotes | 68 | 3 | 4.34E-02 |
| path:mmu05322 | Systemic lupus erythematosus | 29 | 2 | 4.36E-02 |
| path:mmu04610 | Complement and coagulation cascades | 30 | 2 | 4.64E-02 |
| path:mmu05164 | Influenza A | 119 | 4 | 4.84E-02 |
| path:mmu03440 | Homologous recombination | 34 | 2 | 5.81E-02 |
| path:mmu03460 | Fanconi anemia pathway | 44 | 2 | 9.10E-02 |
| path:mmu05171 | Coronavirus disease - COVID-19 | 163 | 4 | 1.19E-01 |
| path:mmu01240 | Biosynthesis of cofactors | 106 | 3 | 1.23E-01 |
| path:mmu05133 | Pertussis | 53 | 2 | 1.24E-01 |
| path:mmu05140 | Leishmaniasis | 54 | 2 | 1.28E-01 |

## Results

### XO4- and FACS-based isolation of Aβ-positive and Aβ-negative microglia

To understand the different effects of ultrasound-mediated BBB opening on plaque-phagocytic and non-phagocytic microglia in AD, we applied SUS^+MB^ or sham (i.e. mice were anaesthetized and injected with microbubbles, but not exposed to ultrasound) to the brains of APP23 mice or WT littermate controls (**Fig. 1A,B**). In addition, to be able to distinguish between microglial cells that had internalized Aβ and those that had not, we used the fluorescent Congo-red derivative methoxy-XO4 to stain Aβ within microglia when injected into live mice, as previously done [9, 25]. This allowed us to use a FACS-based technique to separate and isolate XO4^+^ (Aβ phagocytic) and XO4^-^ (non-phagocytic) microglia following both SUS^+MB^ and sham treatment paradigms (**Fig. 1C,D**).

### Genotype, treatment and Aβ internalization induce distinct phenotypes in microglia

Sorted microglial cells (both XO4^+^ and XO4^-^) from the four experimental groups (**Fig. 1A**) were subjected to RNA isolation, followed by RNA sequencing (RNAseq) and bioinformatic analysis. Principal component analysis (PCA) revealed a clear segregation between the samples of different genotypes, being either of APP23 mutant or WT origin (**Fig. 2A**). In addition, both PCA and hierarchical clustering analysis (**Fig. 2B**) segregated distinct microglial populations induced by Aβ uptake (XO4^+^ versus XO4^-^ cells), as well as treatment (SUS^+MB^ versus sham-treated animals), which were markedly accentuated in the APP23 samples. As the effect of treatment in WT mice was negligible and differential expression analysis had revealed no significant changes to the transcriptome, we subsequently focused our analysis on the effects of ultrasound ± Aβ internalization in APP23-derived microglia only.

### SUS treatment induces an increased number of up-regulated genes in microglia

To gain insight into the response of APP23 microglia to the SUS treatment regime, we further analyzed the transcripts obtained from XO4^+^ and XO4^-^ cells. Our analysis identified 397 differentially enriched genes (FDR≤0.05), with 155 genes being specific for XO4^+^ cells, 199 genes specific for XO4^-^ microglia, and 123 genes being independent of the Aβ signature. Analyzing the treatment-dependency patterns, we observed that most of the up-regulated genes were induced by SUS^+MB^, with a total of 353 enriched genes across all the Aβ internalization levels (**Fig. 3A**), and only 44 genes that were down-regulated following SUS^+MB^ treatment (**Fig. 3B**). These enrichment patterns are highlighted in more detail by the Volcano plots, with both the XO4^+^ cells (**Fig. 4A**) and XO4^-^ cells (**Fig. 4B**) exhibiting increased numbers of differentially enriched genes induced by the SUS^+MB^ treatment. The top 50 differentially expressed genes in the SUS^+MB^ versus sham groups for both XO4^+^ and XO4^-^ microglia are presented in **Tables 1 and 2**.

**Figure 4.**
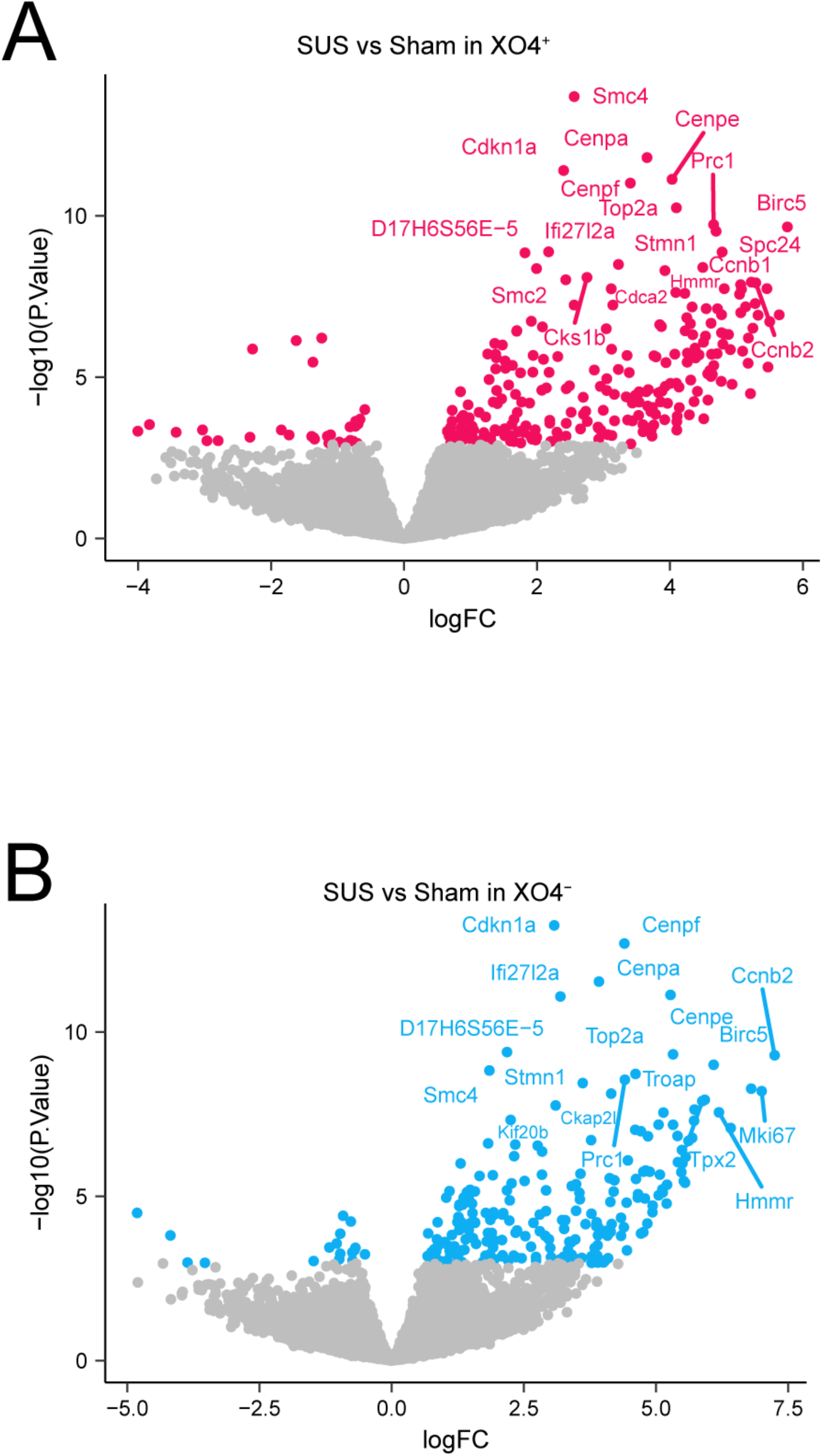
Volcano plots reveal dysregulation of genes in microglia from APP23 mice following SUS^+MB^ or sham treatment. **(A)** XO4^+^ microglia show a large number of genes up-regulated, with the 20 with the largest fold-change (logFC) labeled in magenta. **(B)** XO4^-^ microglia also show a large number of up-regulated genes, with the 20 with the largest fold-change (logFC) labeled in blue.

### SUS treatment induces an enrichment in microglial cell-cycle-related transcriptome

We next sought to identify the functionally enriched pathways induced by SUS treatment and compare them in both the XO4^+^ and XO4^-^ microglia. Applying a gene ontology (GO) enrichment analysis to the SUS^+MB^ versus sham datasets revealed the top 10 enriched pathways that included ‘cell cycle’, ‘DNA replication’ and ‘DNA metabolic processes’ (**Table 3**). KEGG pathway analysis revealed that the most enriched pathways included ‘DNA replication’ and ‘cell cycle’, as well as established pathways in relation to the role of microglia in AD, such as ‘phagosome’ and the ‘complement and coagulation cascade’ (**Table 4**). Inspection of the ‘cell cycle’ and ‘phagosome’ pathways in a treatment-(SUS^+MB^ versus sham) and Aβ internalization (XO4^+^ versus XO4^-^)-dependent manner revealed similar trends, with a stronger response found for the XO4+ microglia containing internalized Aβ (**Fig. 5**). More genes in the phagosome pathway are significantly altered by SUS in XO4+ microglia (7 genes up-regulated and 3 down-regulated) than XO4-microglia (5 genes up-regulated) with 2 of these genes up-regulated in both (**Fig. 5a**). For the cell cycle pathway there were also more genes up-regulated in XO4+ microglia (18 genes up-regulated) than XO4-microglia (11 genes-upregulated with 9 of these genes up-regulated in both (**Fig. 5b**).

**Figure 5.**
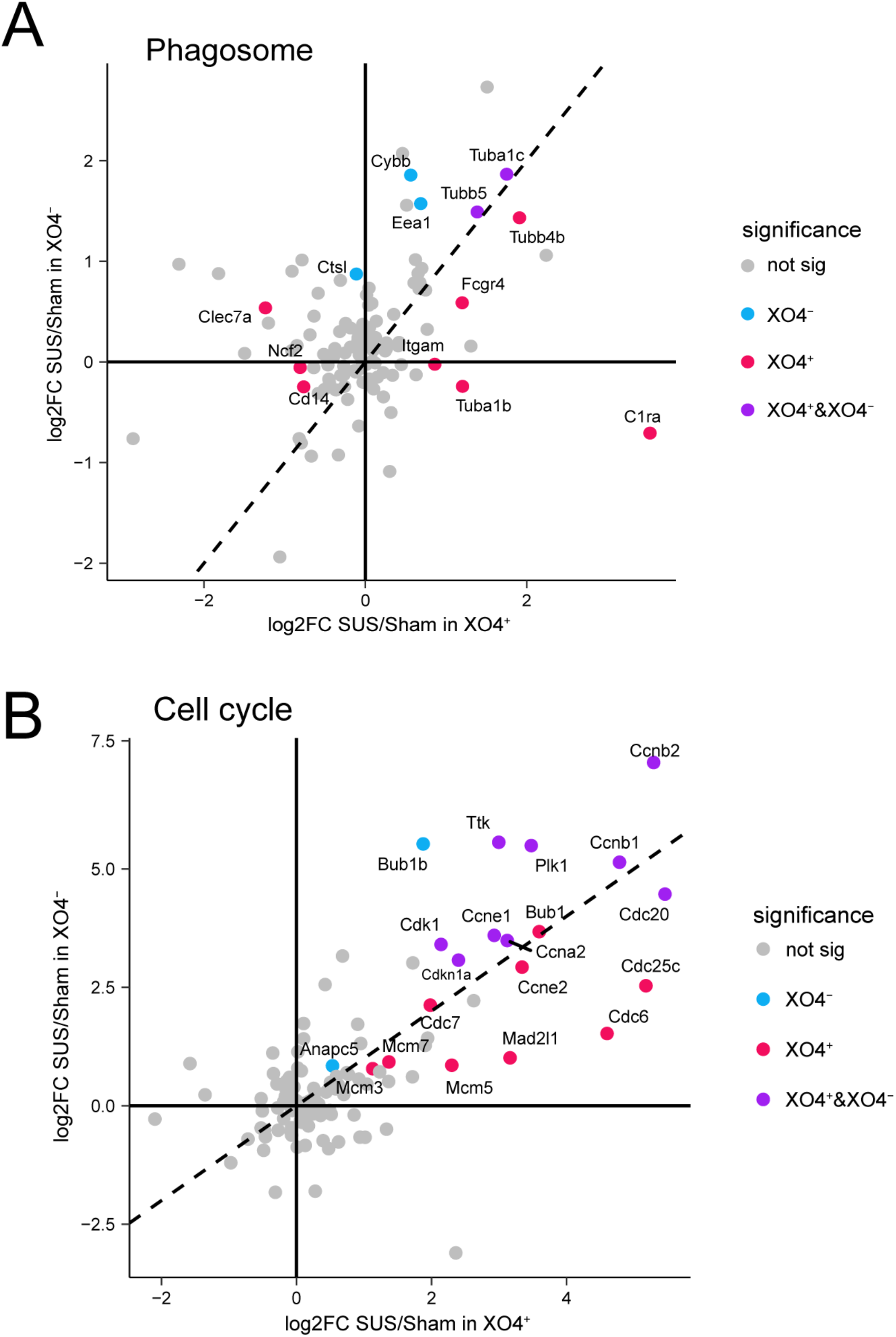
KEGG analysis reveals phagosome and cell cycles as top dysregulated pathways in SUS^+MB^ treated APP23 microglia. **(A)** Plotting fold-changes in gene expression (log2FC) with XO4^+^ (x-axis) and XO4^-^ (y-axis) for phagosome genes indicates that the presence of internalized Aβ (XO4^+^) in microglia treated with SUS^+MB^ leads to an additive effect on this pathway. **(B)** Plotting fold-changes in gene expression (log2FC) with XO4^+^ (x-axis) and XO4^-^ (y-axis) for cell cycle genes reveals that cell cycle genes are upregulated regardless of the presence of internalized Aβ (XO4^+^) in microglia cells, with cell cycle genes being enriched in both XO4^+^ and XO4^-^ cells. Data points are colored according to their statistical significance in the contrasts.

## Discussion

In this study, we sought to investigate how the application of BBB opening achieved with therapeutic ultrasound in conjunction with intravenously injected microbubbles changes the microglial transcriptomic profile in a mouse model of AD. This profile was obtained by analyzing the microglial transcriptome and correlating it with the presence or absence of Aβ in the microglia at 48 hours after treating amyloid-depositing APP23 mice with SUS^+MB^. This allowed us to identify several cellular functions that were increased by SUS^+MB^ application, such as the phagosome (as would have been expected in Aβ-containing XO4^+^ cells), as well as members of the cell cycle pathway, which were upregulated independent of the Aβ internalization level. Our findings further indicate that BBB opening by ultrasound has lasting immunomodulatory effects in the context of AD.

Several previous studies have investigated the effect of ultrasound application to the brain by applying -omics techniques to cell populations. One study investigating ultrasound-mediated delivery of plasmids to the brain of WT mice performed single-cell RNA sequencing and found an upregulation of lysosomal genes in microglia 48 hours after ultrasound treatment [26]. In support of this, in a SWATH quantitative proteomics screen following a series of six weekly sessions of SUS treatments (SUS^+MB^) in aged wild-type C57Bl/6 mice, we identified an increase in two microglial proteins (LRBA and CAGP) which are involved in phagocytosis [27].

In aiming to find a treatment modality for AD, boosting the Aβ phagocytic activity of microglia may present a promising strategy if applied to the earliest stages of the disease, by increasing the clearance of protein deposits [28]. A previous attempt to investigate the microglial response following ultrasound treatment focused on investigating transcripts related to the downstream effects of the NFκB pathway and damage-associated molecules in bulk lysates from WT rodent brains, with most transcript levels returning to baseline after 24 hours [29]. The study concluded that ultrasound treatment with BBB opening may lead to sterile inflammation. A subsequent study, however, reported no significant changes in the expression of any of the NFκB-related genes when using a lower, more clinically relevant dose of microbubbles [30]. These opposing effects could be attributed to the specific ultrasound treatment parameters that elicit a cavitation-modulated inflammatory response through the microbubbles present in the blood circulation [31]. In addition, the transcriptomic response to ultrasound-induced BBB opening was found to be dependent on the type of anesthesia used during the procedure [26]. Of note, we used ultrasound settings that we have previously demonstrated to increase microglial phagocytosis [13], cause no damage to neurons [32], and which we suggest have relatively lower levels of cavitation and BBB opening. Taken together, these results indicate that the magnitude of the effects of ultrasound-mediated BBB opening are heavily dependent on the parameters of ultrasound used as well as the downstream analysis (RNA or protein, and heterogeneous tissue or isolated cell-types).

Using a protocol that we have recently applied to reveal the transcriptional signature of microglia associated with Aβ phagocytosis [9], in the current study, we aimed to investigate changes after SUS treatment in the APP23 model of AD. Our results revealed enriched pathways in the Aβ-containing microglia, such as cell cycle, phagosome, complement activity, and metabolism. Some of the pathways identified by our analysis have previously been associated with microglial activation in AD, validating our results. Indeed, microglia have been observed to remove synapses in AD through a mechanism involving members of the complement system [33, 34]. In addition, it has been proposed that the metabolism of microglia is impaired in AD, an effect that can be ameliorated by enhancing the cellular energetic and biosynthetic metabolism [35]. Increased microglia numbers in proximity to plaques are associated with more compact plaques and reduced axonal dystrophy [36], and we have previously reported increased microglial numbers around plaques following SUS^+MB^ treatment [17]. Higher numbers of microglia around plaques may result from an increased proliferation or metabolic activity, as hinted in the present study. The reactivation of the cell-cycle machinery in microglia following ultrasound treatment is of particular interest, as it has recently been reported that repopulating microglial cells following ablation are neuroprotective in AD [37, 38].

In conclusion, we have examined the effect of SUS^+MB^ on the microglial transcriptome in the presence or absence of amyloid pathology. SUS^+MB^ leads to temporary opening of the BBB and alters microglial gene expression in the AD brain to modulate several cellular pathways, including the cell cycle, various metabolic pathways and the phagosome. Harnessing the protective effects of microglia in the context of AD could potentially be achieved by a combination of SUS^+^MB-mediated BBB opening and targeted drug delivery.

## Abbreviations

AD: Alzheimer’s disease
Aβ: amyloid-beta
BBB: blood brain barrier
FACS: fluorescence activated cell sorting
MBs: microbubbles
NFκB: nuclear factor kappa light chain enhancer of activated B cells
PBS: phosphate-buffered-saline
PCA: principal components analysis
SUS: scanning ultrasound
WT: wild-type

## Credit author statement

GL, GS and AG performed experiments. GL, LGB, JS, YC, and AG analyzed data. GL, LGB, JS, JP and JG wrote the manuscript with input from all the authors.

## Acknowledgements

The authors would like to acknowledge Flowcore, Monash Health Translation Precinct Medical Genomics Facility, and Australian Research Laboratories/Monash Animal Research Platform, Monash University, for the provision of instrumentation, training and technical support. We thank Tishila Palliyaguru and Linda Cumner for assistance with animal colonies, and Rowan Tweedale for critical reading of the manuscript.

